# A regulatory-sequence classifier with a neural network for genomic information processing

**DOI:** 10.1101/355974

**Authors:** Koh Onimaru, Osamu Nishimura, Shigehiro Kuraku

## Abstract

Genotype-phenotype mapping is one of the fundamental challenges in biology. The difficulties stem in part from the large amount of sequence information and the puzzling genomic code, particularly of non-protein-coding regions such as gene regulatory sequences. However, recently deep learning–based methods were shown to have the ability to decipher the gene regulatory code of genomes. Still, prediction accuracy needs improvement. Here, we report the design of convolution layers that efficiently process genomic sequence information and developed a software, DeepGMAP, to train and compare different deep learning-based models (https://github.com/koonimaru/DeepGMAP). First, we demonstrate that our convolution layers, termed forward- and reverse-sequence scan (FRSS) layers, enhance the power to predict gene regulatory sequences. Second, we assessed previous studies and identified problems associated with data structures that caused overfitting. Finally, we introduce several visualization methods that provide insights into the syntax of gene regulatory sequences.

## Introduction

In the last decade, advances in DNA sequencing technologies have dramatically increased the amount of genome sequence data derived from diverse species (1) as well as from individual humans (2). The next demanding challenge is the deeper understanding of how genome sequences encode organismal traits and how functional information can be extracted (3). Such sequence-based understanding would ultimately enable the prediction of phenotypes based on genome sequence information, i.e., genotype-phenotype mapping. The syntax of protein-coding genes is well understood, e.g., certain phenotypic consequences are predictable (such as nonsense mutations), yet the basic rules for non-coding sequences have not been completely established. Several projects including ENCODE (4,5), ROADMAP (6), and FANTOM (7) have accumulated epigenomic data to annotate the characteristics of non-coding sequences, but a comprehensive understanding of the association between epigenomic signatures and sequence information has yet to be attained.

Recently, this challenge has been addressed with deep learning-based methods (8–11). Deep learning is a subfield of machine-learning methods, and has been applied to a variety of problems such as image classification and speech recognition (12). Although much research has used deep-learning methods to automate routine but complex tasks that only humans were previously able to carry out, some cases are intended to surpass the human ability, such as playing the game “Go”(13). For the application of deep learning to genomics, convolutional neural networks (CNN) or recurrent neural networks (RNN) are trained with input genomic sequences that are labeled with epigenomic data to predict functional noncoding sequences. Once trained, these types of classifiers can infer the functional effect of mutations in non-coding genomic regions from individual genome sequences. However, even though these deep learning–based classifications have outperformed other methods, such as the support vector machine (14), the accuracy of prediction is still far from satisfactory. the prediction accuracy is still far from satisfactory. In addition, as it is generally known, it remains elusive as to what deep learning-based models actually “learn” and what kinds of “understanding” underlies their predictions (15). In the present study, we first describe a CNN-based classifier that can outperform state-of-art models. Second, we identified problems associated with data structures that caused overfitting. Finally, we introduce methods to visualize trained models, potentially revealing the general syntax underlying regulatory sequences.

## Results

To improve the accuracy of predicting regulatory sequences, we devised a deep learning-based method with three main features: a) integrating information from forward and reverse DNA sequences; b) simplifying the data structure; c) a new quality index to filter out low-quality data from a training dataset.

As training data, we downloaded several alignment files containing data from chromatin accessibility assays and chromatin immunoprecipitation-sequencing (ChIP-seq) experiments from the ENCODE project, and regions enriched with reads were determined as peaks by MACS2 peak caller (16). These data were used to mark genome sequences. We divided each of the mouse and human genome sequences into 1000-basepair (bp) windows and converted the four letters (i.e., A, C, G, T) into one-hot vectors (four dimensional zero-one vectors). To denote epigenomic marks, we assigned each window as 1 if it overlapped a signal positive region or 0 otherwise (signal negative) (see Methods for details). To reduce the number of potential artifacts caused by window boundaries, we also added windows that were shifted by 500 bp toward the 3’ side (Fig. 1a; we refer to this window structure as 1-kbp window/0.5-kbp stride). These data were used to train CNN models (see Supplementary Fig. 1 for the overall scheme).

**Fig 1.**
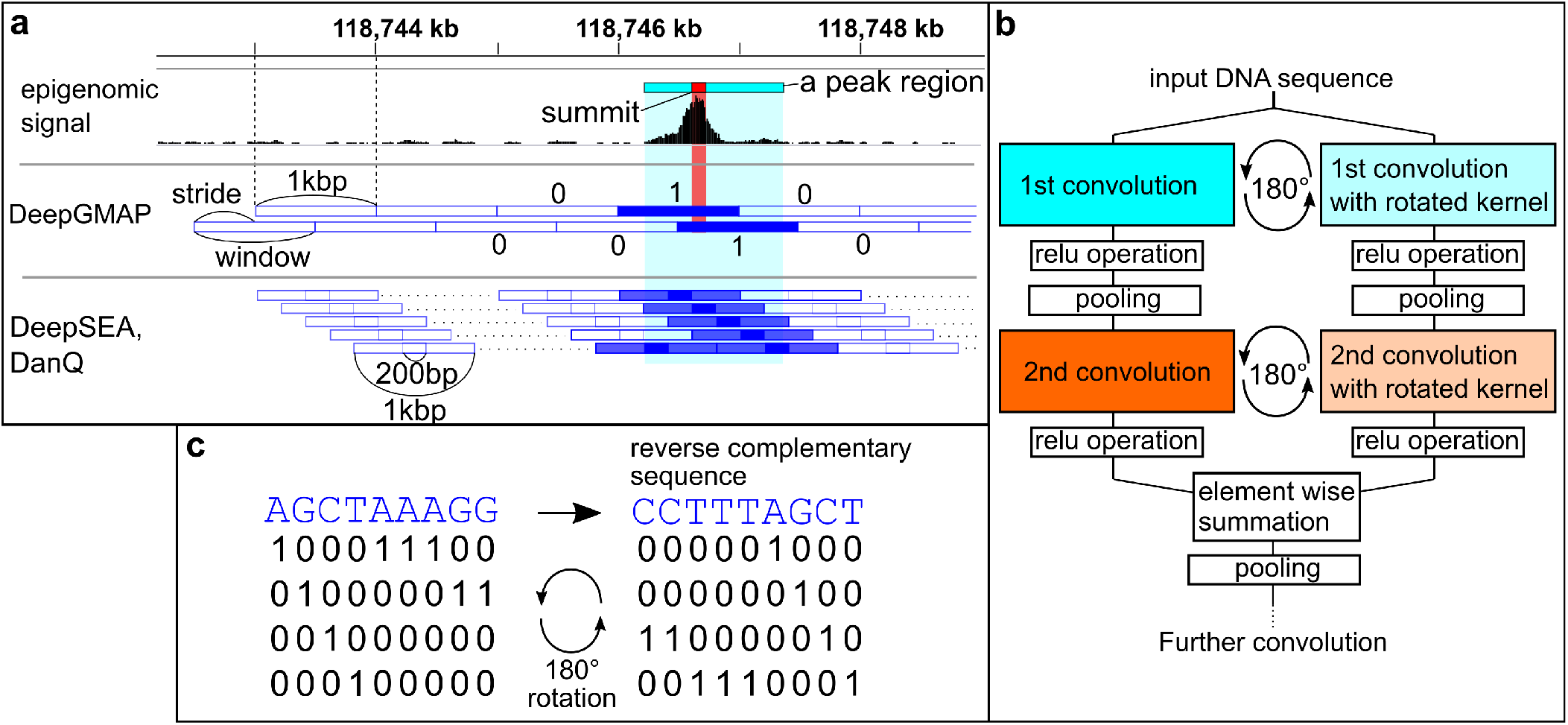
The forward- and reverse-sequence scan (FRSS) layers and data structures. **a**, Subdivision of genome sequences into windows, and assignment of epigenomic signals. A peak region (red rectangle) is determined from epigenomic data and assigned as “1” to overlapping genomic regions (blue-filled rectangle), otherwise “0” (blue, empty rectangle). DeepGMAP, this study; DeepSEA, DanQ, previous studies. **b**, The architecture of the FRSS layers. An input sequence is scanned by the first convolution kernels, and the kernels are rotated in parallel. After implementing the relu operation, pooling, and the second convolution, the two parallel outputs are combined through summation. c, Illustration of the 180-degree rotation of kernels. In this example, a 9×4 kernel is trained to detect a sequence, AGCTAAAG (left). The geometric rotation of the kernel (right) results in the reverse complement of the original sequence.

To find an effective architecture of neural networks, we first compared the performance of published models such as CNN-based models (DeepSEA and Basset) and a CNN-RNN hybrid (DanQ; see Supplementary Table 1 for details of the models). As a quick benchmark, we trained them with a subset of mouse DNase-seq data (Supplementary Table 2) because DNase-seq data generally include diverse regulatory elements (17). To reduce computation time, we limited the number of training epochs to one, which means that models were trained with the full dataset only once. For evaluation, we used the entire chromosome 2 sequence, which was excluded from training datasets, and calculated the prediction accuracy of models using two scores, namely the area under the receiver-operation curve (AUROC) and the area under the precision-recall curve (AUPRC). Because AUROC is not a proper criterion if the majority of the data is negatively labeled, we considered AUPRC as a more important criterion (18). Consequently, the DeepSEA architecture performed better than Basset and DanQ (Table 1). The relatively poor performance of DanQ seemed inconsistent with a previous report (10), and this was probably attributable to an insufficient amount of training, i.e., the DanQ model requires 60 epochs to attain good performance (10).

**Table 1.** A performance comparison between models

|  | 129_ES_E14 |  | forebrain E11.5 |  | forelimb E11.5 |  | mean |  | time<sup>b</sup> |
| --- | --- | --- | --- | --- | --- | --- | --- | --- | --- |
|  | AUROC | AUPRC | AUROC | AUPRC | AUROC | AUPRC | AUROC | AUPRC |  |
| DeepSEA | 0.9234 | 0.6038 | 0.9141 | 0.6380 | 0.9315 | 0.5734 | 0.9230 | 0.6051 | 0.1773 |
| Basset | 0.9085 | 0.5796 | 0.9107 | 0.6280 | 0.9175 | 0.5476 | 0.9123 | 0.5851 | 0.0847 |
| DanQ | 0.9057 | 0.5785 | 0.9076 | 0.6214 | 0.9046 | 0.5164 | 0.9059 | 0.5721 | 0.4586 |
| DanQ block<sup>a</sup> | 0.9053 | 0.5699 | 0.9063 | 0.6168 | 0.9030 | 0.5031 | 0.9049 | 0.5633 | 0.4832 |
| conv4 | 0.9300 | 0.6234 | 0.9146 | 0.6426 | 0.9351 | 0.5884 | 0.9266 | 0.6181 | 0.2548 |
| conv3-FRSS | 0.9364 | 0.6447 | 0.9171 | 0.6496 | 0.9421 | 0.6098 | 0.9319 | 0.6347 | 0.3636 |
| conv4-FRSS | **0.9414** | **0.6571** | **0.9178** | **0.6520** | **0.9476** | **0.6280** | **0.9356** | **0.6457** | 0.3729 |
| conv4-FRSS+ | **0.9418** | **0.6631** | **0.9183** | **0.6549** | **0.9475** | **0.6347** | **0.9359** | **0.6509** | 0.5802 |
| DanQ-FRSS | 0.9365 | 0.6523 | 0.9162 | 0.6485 | 0.9453 | 0.6222 | 0.9327 | 0.6410 | 0.7905 |

<sup>a</sup>Tensorflow has several LSTM cells, and this model uses LSTMBlockCell, and the above uses LSTMCell. <sup>b</sup>the mean time (second) of 20 minibatch updates.

Based on this result, we next explored whether better models could be attained by modifying the DeepSEA architecture. First, as more stratified convolutional networks are known to perform better (19), we inserted an additional convolution layer between the third convolution layer and the first fully connected layer of the DeepSEA model (with a reduced kernel number in some layers and smaller pooling patches). As expected, the four-layer convolutions achieved slightly better accuracy (conv4 in Table 1). We next focused on one of the fundamental characteristics of genome sequences—the forward and reverse strands. In previous studies, the information on reverse-strand sequences was ignored or used as independent data. However, for example, transcription factors with leucine-zipper or helix-loop-helix domains bind both strands through dimerization, and such binding thus results in palindromic binding motifs (20,21). Inspired by this fact, we devised a set of layers that can integrate information from the both strands (Fig. 1b and Methods). We arranged the one-hot vectors that represent AGCT in a symmetric manner so that a 180-degree rotation of the one-hot vectors results in a reverse complementary sequence (Fig. 1c). Thus, reverse-sequence information can be processed by rotating the kernels 180 degrees. The hidden layers derived from the two strands are combined by element-wise addition after the second convolution. We termed this architecture FRSS (forward- and reverse-sequence scan) layers. The replacement of the first two convolutional layers in the DeepSEA model with FRSS (conv3-FRSS in Table 1) improved the accuracy of prediction more than the simple addition of a convolution layer (conv4). Moreover, a four-layer convolution model with FRSS achieved higher AUPRC scores than the other models (conv4-FRSS in Table 1, and see Supplementary Fig. 2 for a visual comparison). Raising the kernel numbers of each convolution layer also slightly improved the performance (conv4-FRSS+ in Table 1) but also increased computation time (the right most column in Table 1). We also found that the FRSS layers increased the learning efficiency of the DanQ model by a level comparable with conv4-FRSS (DanQ-FRSS in Table 1). These results were consistently reproduced by repeated training and testing (Supplementary Table 3), indicating that the benchmark is robust against random initialization. Together, these results showed that the FRSS layers enhance the predictive power of deep learning-based models.

Next, we examined the structure and quality of training data using the conv4-FRSS model. In certain previous studies, epigenomic signals were distributed into 200-bp windows of genome sequences, and 400-bp extra-sequences were added to the left and right sides of each window (9) (referred to as “DeepSEA-type data” in this study; Fig. 1a). For comparison, we trained conv4-FRSS with the DeepSEA-type data structure. Training with the DeepSEA-type data yielded a higher AUROC score but a lower AUPRC score (DeepSEA-type in Table 2). The high value of the AUROC is probably attributable to an increase in negative labels. In addition, we tested several window and stride sizes and found that models can be trained most efficiently with a 1-kbp window and 0.3-kbp stride data. As shown in Table 2, training with small-sized strides such as 0.1 and 0.2 kbp (and also DeepSEA-type data) yielded higher AUPRC scores when tested with the chromosome 1 sequence, which was included in the training data, than tested with the test dataset (chromosome 2, which was excluded from the training data), indicating overfitting to redundant sequences in the training dataset. This result also explains the unusually high AUPRC scores reported by Min et al. (2017) (22), which is inconsistent with the results of Zhou & Troyanskaya (2015) (9)). Because Min et al. trained CNN-based classifiers with 2-bp stride data and evaluated their model by resampling training data, the high AUPRCs in their report seemed to result from overfitting. We also examined peak callers. Although we utilized MACS2 to determine peak regions, previous studies used peak regions provided by the ENCODE project, which utilized Hotspot (23) as the peak caller. As shown in Table 2, training with Hotspot peaks yielded lower prediction accuracy (in Table 2). These results indicated that data-structure design is a critical factor for training models efficiently and that data augmentation using small strides, which was implemented in several studies (22,24), is not an optimal strategy for training models.

**Table 2.**
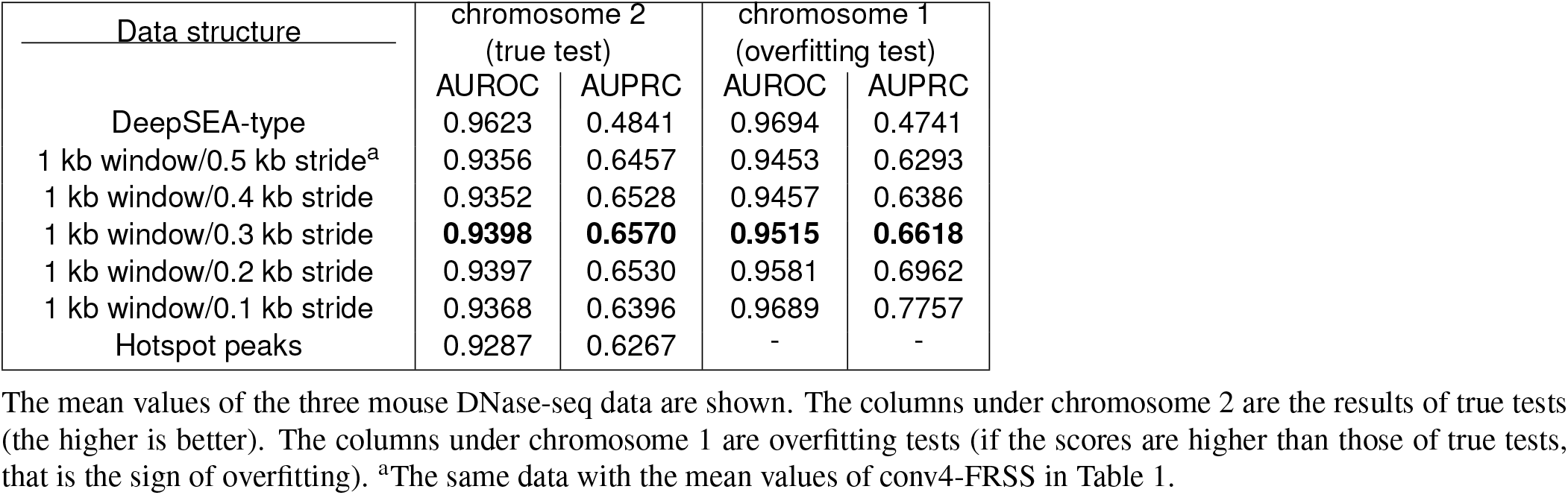
A performance comparison between data structures

| Data structure | chromosome 2<br>(true test) |  | chromosome 1<br>(overfitting test) |  |
| --- | --- | --- | --- | --- |
|  | AUROC | AUPRC | AUROC | AUPRC |
| DeepSEA-type | 0.9623 | 0.4841 | 0.9694 | 0.4741 |
| 1 kb window/0.5 kb stride<sup>a</sup> | 0.9356 | 0.6457 | 0.9453 | 0.6293 |
| 1 kb window/0.4 kb stride | 0.9352 | 0.6528 | 0.9457 | 0.6386 |
| 1 kb window/0.3 kb stride | **0.9398** | **0.6570** | **0.9515** | **0.6618** |
| 1 kb window/0.2 kb stride | 0.9397 | 0.6530 | 0.9581 | 0.6962 |
| 1 kb window/0.1 kb stride | 0.9368 | 0.6396 | 0.9689 | 0.7757 |
| Hotspot peaks | 0.9287 | 0.6267 | - | - |

To validate the aforementioned quick benchmark, we extended the list of training data. First, we newly trained conv4-FRSS with the same set of 125 human DNase-seq data used by Zhou & Troyanskaya (2015) (9) (see Supplementary Table 2 for the lists of data). We found that conv4-FRSS significantly outperformed DeepSEA and DanQ (Fig. 2a and Supplementary Table 4; the scores of DeepSEA and DanQ were obtained from their publications). We also trained conv4-FRSS with 365 and 93 of human and mouse DNase-seq peaks, respectively, generated by MACS2 (see Supplementary Table 2 for the lists of data). As shown in Fig. 2a and Supplementary Table 4, we noticed that training with mouse data yielded higher performance than with human data. The cause may be attributable to a difference in genome size, genetic variation (should be lower for inbred mice), or species-specific trends such as gene density and GC content. We also focused on CTCF ChIP-seq data because (a) CTCF has a general function in genome organization, (b) ENCODE releases plenty of CTCF data for both mice and humans, and (c) as evident for the AUPRC scores of DeepSEA (Supplementary Fig. 3; reanalyzed data from Zhou & Troyanskaya, 2015 (9)), the prediction accuracy is highly dependent on the target of ChIP-seq, and that of CTCF is the highest among transcription factors. As a result, the conv4-FRSS model was efficiently trained with both human and mouse CTCF data, and this model yielded a median AUPRC score of >0.70 (see Fig. 2b and 2c for a visual comparison between predictions and original signals). A comparison between the predictions of the model and CTCF motif regions detected by FIMO (25) [CTCF motif (FIMO) in Fig. 2c] revealed that the model can predict CTCF binding regions that do not contain the canonical CTCF motif. In addition, we also used the class saliency extraction method (26) to identify informative sequences at the single-nucleotide level (Supplementary Fig. 4). This method is equivalent to the in silico saturation mutagenesis approach used previously (8,9) but is more computationally efficient because it does not require mutation-by-mutation evaluation (see Methods for details). Furthermore, using the dog genome (CanFam3.1), we performed a crossspecies prediction of CTCF binding regions with the model trained solely with mouse data. As shown in the bottom panel of Fig. 2c, the predictions of the model matched with the real CTCF signals of dog liver (27) and detected species-specific differences (red rectangles in Fig. 2c), indicating the generality of the model. Based on all these data, we concluded that conv4-FRSS can be applied to diverse data.

In the above training with extended dataset, we suspected that several classes with low AUPRCs were attributable to poor data quality. To address this possibility, we calculated a known quality index, FRiP (fraction of reads in peaks (28)) of mouse CTCF data and found that classes with low AUPRCs tended to have low FRiPs (Fig. 2d), with some exceptions (arrows in Fig. 2d). Because FRiP is calculated based on reads in peaks per total reads, values may be inflated when the read number of the source data is too low. To correct such inflation, we multiplied FRiP by the fraction of genomic regions covered by at least one read in a genome. As shown in Fig. 2d, the corrected FRiPs (cFRiPs) could distinguish high and low AUPRC data more clearly. In addition, cFRiP yielded a stronger correlation with the total peak numbers of data than the uncorrected FRiP (Fig. 2e and Supplementary Table 5). Although the other datasets showed weaker relations between FRiP and AUPRC, data with optimal cutoff values of cFRiP increased the average AUPRC score except for the human CTCF data (filtered hg38 and mm10 data in Fig. 2a and 2b, and Supplementary Fig. 5). These results suggested two points. First, the correction method reasonably represents the true quality of the data. Second, the linear correlation between peak numbers and cFRiPs implies that data quality is not saturated enough to detect all true peaks (and hence, the possibility remains that the model learns sample quality rather than cell-specific patterns). Therefore, whereas the ENCODE project seems to consider ≥1% FRiP as good data (28), our results suggest that higher stringency during quality control is required for the precise annotation of genomes.

**Fig 2.**
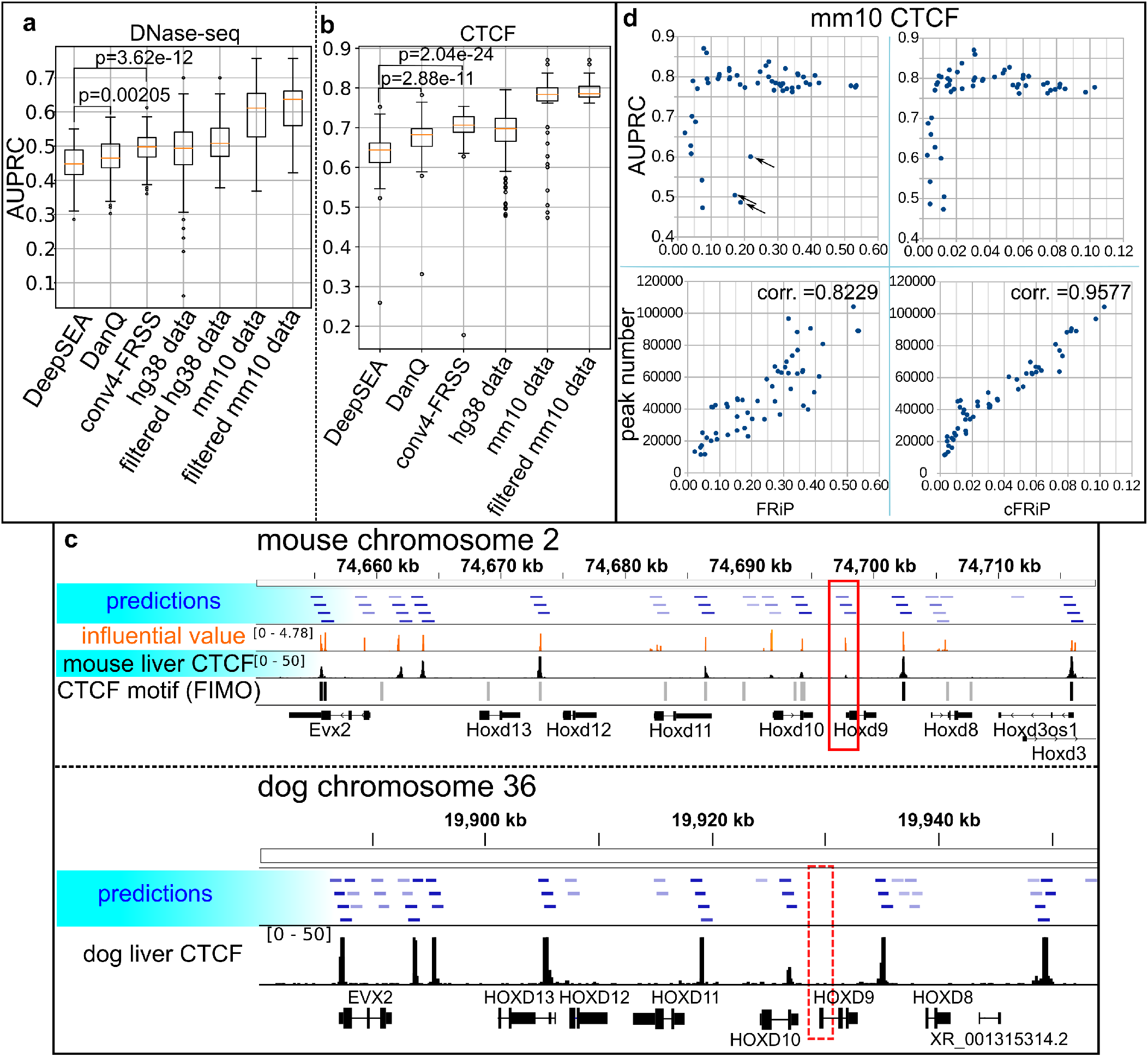
Performance analyses of models with extended data sets. **a, b**, Comparison of AUPRCs of DNase-seq data (a) and CTCF data (b). DeepSEA and DanQ, AUPRCs obtained from their original reports; conv4-FRSS, predictions trained with the same peak files, but a different data structure (1-kbp window/0.3-kbp stride); hg38 and mm10 data, 365 and 93 DNase-seq datasets generated by the MACS2 peak caller; filtered hg38 and mm10 data, DNase-seq data with high cFRiPs. p, two-sided Wilcoxon-Mann-Whitney test. Boxes, the lower to upper quantile values of the data; orange lines, the median; whiskers, the range of the data (Q1 – IQR × 1.5 and Q3 + IQR × 1.5 for the lower and upper bounds, respectively); flier points, those past the end of the whiskers. **c**, An example of CTCF binding site predictions on the HoxD cluster of mice and dogs. predictions, CTCF binding regions predicted by conv4-FRSS (only with prediction values >= 0.2 are shown, and the bluer is more probable); influential value, values calculated with the class saliency extraction method; mouse/dog liver CTCF, alignment data from ENCFF627DYN and ERR022304, respectively; CTCF motif (FIMO), sites that were detected by FIMO (black boxes, p <= 1e-4; gray boxes, 1e-4 < p <= 1e-3); red rectangles, a CTCF peak that is present in the mouse genome (solid rectangle) but not the dog genome (dashed rectangle). d, Scatter plots for FRiP and corrected FRiP with AUPRCs (top panels) and peak numbers (bottom panels). corr., Spearman correlation.

To obtain insights into how the models predict regulatory sequences and their syntax, we visualized the model (conv4-FRSS) trained with the subset of mouse DNase-seq data. First, as has been done in previous studies (8,10),we analyzed individual kernels of the first layer, which directly interact with DNA sequences and thus represent DNA motifs that are important for classifying regulatory sequences. We converted the weight variables to probability matrices by using a softmax-like operation and found that established models have indeed learned many known transcription factor binding motifs, such as those bound by CTCF, Sox9, OCT4 and KLF4 (Fig. 3a, and Supplementary Fig. 6). Because gene regulation often requires the participation of several transcription factors (29), we visualized linkages between kernels and layers (Fig. 3b is linked to the 129 ES E14 class of the three DNase-seq data as an example, and Supplementary Fig. 7 for larger images of linkages to three classes). A comprehensive visualization of linkages is impossible owing to the large numbers of kernels and variables. Therefore, we calculated the strength of linkages by summing weight variables for each kernel, and top 10% of linkages to each class is shown in Fig. 3b and Supplementary Fig. 7. This partial visualization revealed that the information from the first layer (“hidden1”) was gathered at relatively few nodes of the last convolution layer (“hidden4”). We also found that most of the strongly contributing kernels were shared between three classes. In addition, even if a kernel was found to be shared between two classes, the linkages to the last layers differed (kernels with a light blue rectangle and red colored linkages in Supplementary Fig. 7), suggesting that different combinations of kernels were used for the class-specific predictions. Another interesting point is the temporal development of kernels and connections. As shown in Supplementary Fig. 8, the model seems to learn GC-rich sequences (probably recognizing CpG islands or the CTCF binding motif) in early training stages, and kernels are connected by relatively few nodes of the second layer (“hidden2”). In later stages, the model increases the variation and connection of kernels. Taken together, these visualizations suggested that the model seems to discern regulatory sequences via analysis of a combination of kernels.

**Fig 3.**
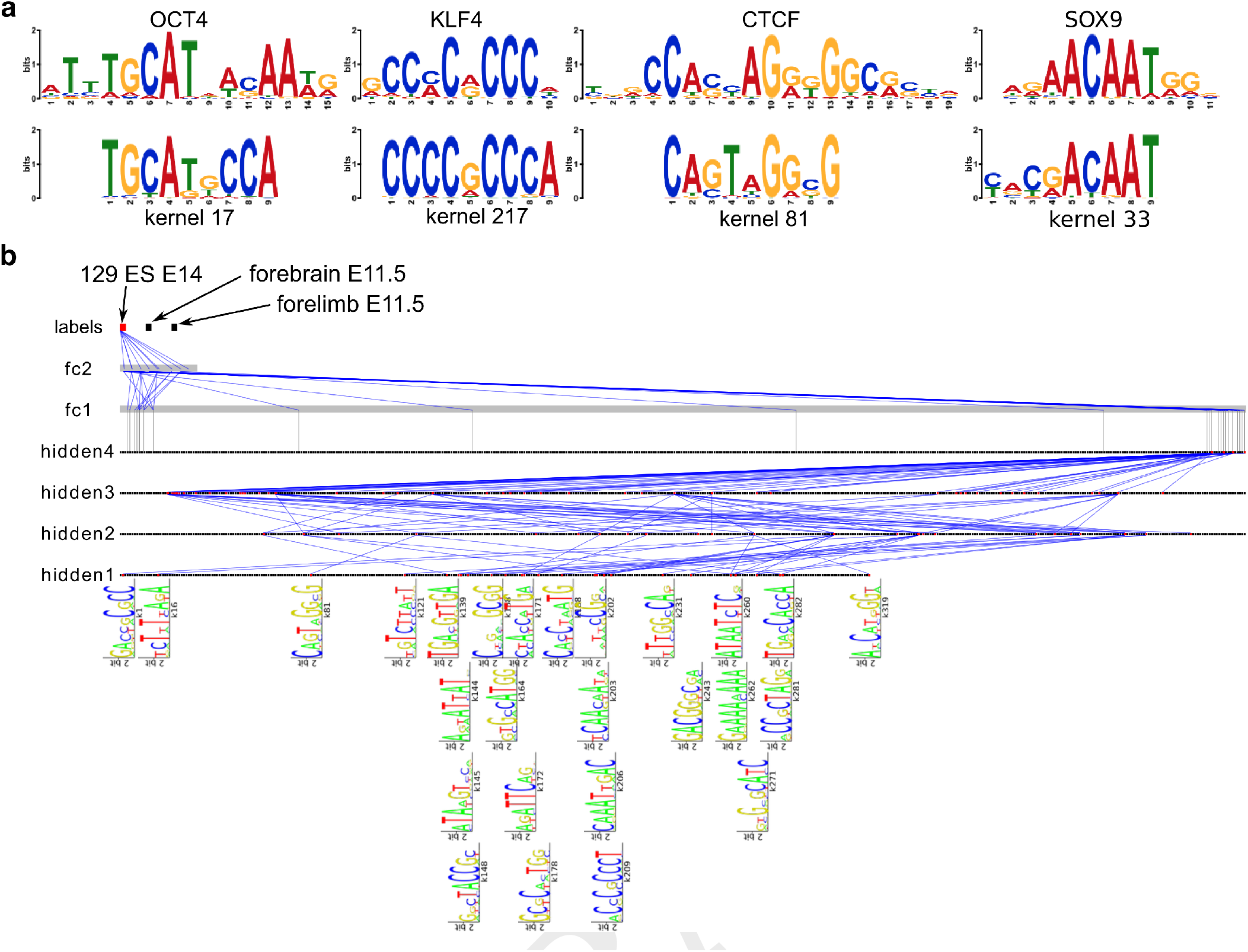
Visualization of trained kernels and their connections. **a**, Examples of known motifs (top) that match with kernels. **b**, Connections of kernels and layers of conv4 with FRSS. As an example, connections that are linked to the 129 ES E14 DNase-seq class are shown. Dots of hidden1-4, the outputs of convolution operations of kernels; fc1 and fc2, the fully connected layers (because there were too many neurons, they are represented as gray bars); labels, the last output layer; blue lines, connections between kernels and layers. Larger images are available in Supplementary Fig. 6.

Next, we applied the activation maximization method25,29,30, in which we look for DNA sequences that maximize neuron activities in the final layer of a trained model by training the sequence itself (see Methods for formal mathematical expressions). First, we trained a sequence to activate the 129 ES E14-specific neuron in the last output layer. As with image-recognition studies (26,30,31), this method generated randomly repeated sequences that probably capture the 129 ES E14 class-specific traits (left panel in Fig. 4). Using the motif comparison software Tomtom (32), we found that the generated sequence contained several OCT4/SOX2 and KLF4 motifs, which are important for the pluripotency of embryonic stem cells (33). For comparison, we also trained a sequence to activate all of the three classes of the neurons, resulting in a CTCF motif–rich sequence (right panel in Fig. 4). This result is consistent with the general role of CTCF as an enhancer looping factor (34). In addition, the generated sequence also contained small motifs next to the well-known 20-bp motif (dashed boxes in the right panel of Fig. 4), which resembled the other part of the alternative long CTCF motif (33/34 bp in length, including an additional GGNANTGCA or TGCANTNCC sequence) (27). Thus, the detection of motifs that are longer than the kernel size constitutes one of the advantages of this activation maximization method. Taken together, these visualization methods would be useful for understanding how deep-learning models predict enhancer sequences.

**Fig 4.**
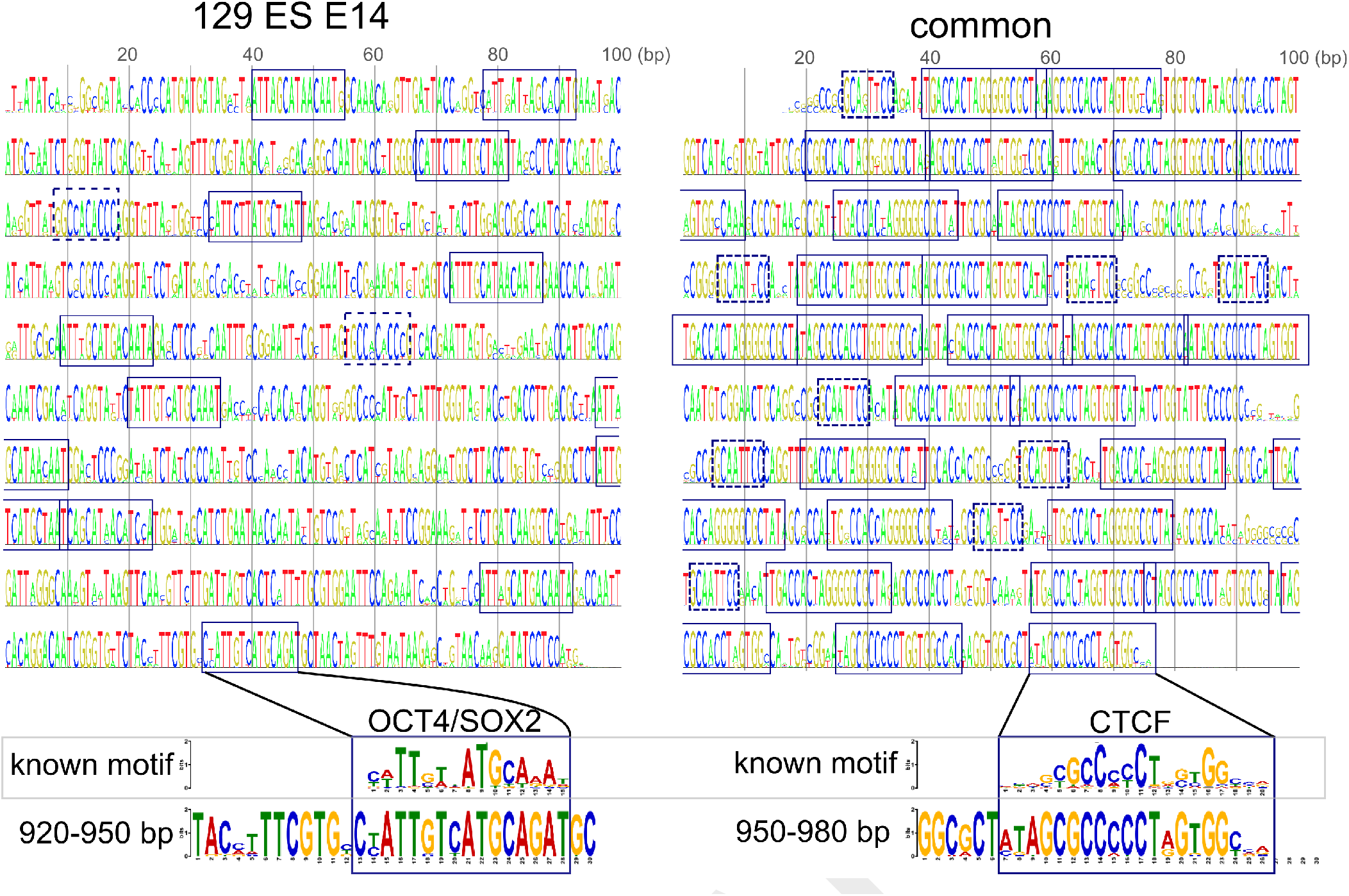
Results of the activation maximization method. Sequences trained to activate the 129 ES E14-specific neuron in the last layer (left panel) and all neurons in the last layer (right panel), respectively. Boxes in the left panel, OCT4/SOX2 binding motifs; dashed boxes in the left panel, KLF4 binding motifs; boxes in the right panel, CTCF binding motifs; dashed boxes, motifs similar to the second part of the alternative longer CTCF motif.

## Discussion

Our results demonstrate that a new convolution architecture, FRSS, efficiently discerns patterns of regulatory DNA sequences. Furthermore, the analyses of data quality and structure revealed that training efficiencies highly depend on the means by which the source data are filtered and processed. Although the visualization methods are still in development, these methods will provide researchers with clues for understanding the syntax of gene regulatory sequences.

We designed FRSS to process the information for forward and reverse sequences simultaneously, and we demonstrated the efficacy of FRSS for this purpose. Contrary to other studies, this is the first to devise a new CNN architecture specialized for processing DNA sequences. However, there are many ways for designing layers for this purpose. For example, instead of pair-wise summation in the last layer of FRSS, one can concatenate the output tensors similar to bidirectional RNN (35). In addition, during finishing this study, we found a study that implemented a CNN model with a similar idea but a different architecture, although their model seemed to suffer from a run-time memory problem, and they did not clearly compare the performance of their model with that of other models (36). We choose the current architecture because its computational cost is smaller than that of other tested architectures, better architectures may be designed that retain low computational cost yet offer greater prediction power. In conclusion, our study constitutes a new paradigm for developing deep learning–based sequence classifiers.

## Methods

### Dataset and processing

The sequences for the human (hg38) and mouse (mm10) genomes were downloaded from UCSC (http://hgdownload.soe.ucsc.edu/goldenPath/). Mitochondrial DNA sequences were excluded from analyses. The genomic sequences were divided into 1-kbp windows with a variety of strides as described in Results. The filtered alignment files for epigenomic data were downloaded from the ENCODE website (https://www.encodeproject.org/; see Supplementary Table 2 for details). A peak caller, MACS2 version 2.1.1.20160309, was used to determine signal peak regions with the following options: “callpeak --call-summits -t <target bam file> -c <control bam file> -f <BAM or BAMPE> -g <hs or mm> -q 0.01” for ChIP-seq, and “callpeak --call-summits -q 0.01 --nomodel --shift -100 --extsize 200 -t <target bam file> -f <BAM or BAMPE> -g <hs or mm>“ for DNase-seq. Using the outputs of MACS2, bedtools (37), and our codes, genomic regions were designated as positive if a window was found to overlap with the summit of a peak. Data for which there was less than 10,000 peaks were excluded from training data. For comparative analyses in Fig. 2a and 2b, peak files were downloaded from http://hgdownload.cse.ucsc.edu/goldenPath/hg19/encodeDCC/. For cross-species comparison, CanFam 3.1 was downloaded from NCBI (https://www.ncbi.nlm.nih.gov/genome/?term=dog), and the short reads for CTCF ChIP-seq data (ERR022304) were downloaded from EMBL-EBI ArrayExpress (https://www.ebi.ac.uk/arrayexpress/experiments/E-MTAB-437/).

The DNA symbols, A, G, C, T and N were converted into one-hot vectors: (1, 0, 0, 0), (0, 1, 0, 0), (0, 0, 1, 0), (0, 0, 0, 1), and (0, 0, 0, 0), respectively. Therefore, one training sample became a 1000×4 tensor. With a mini-batch number of 100 for the stochastic gradient descent and channel dimension 1 (a dimension for colors if the input was image data), the final shape of the input tensor was 100×1000×4×1, with a label tensor of size mini-batch number×class number [i.e., 100 label vectors, and each label looks like (0, 0, 0, 1, 0, 1, …., 0); see Supplementary Fig. 1 for visual illustration].

### Design of models and implementation

Tensorflow r1.8 for python (https://www.tensorflow.org/) with CUDA Driver Version 9.0 and cuDNN v7.0.5 was used as the main machine-learning library for model implementation. The Tensorflow library was compiled from the source codes with the bazel compiler (https://bazel.build/; with options: “c opt --copt=-mavx --copt=-mavx2 --copt=-mfma --copt=-mfpmath=both --copt=-msse4.2 --config=cuda -k //tensorflow/tools/pip_package:build_pip_package”). For convolutions, we used a module, tensorflow.nn.conv2d, with options: “strides=[1, 1, 1, 1], padding=VALID”. For the Basset model, because batch normalization did not yield any positive effects, we removed this operation. For RNN layers, we used the long short-term memory cells (38) (tensorflow.nn.rnn_cell.LSTMCell or tensor-flow.contrib.rnn.LSTMBlockCell without the peephole option), and tensorflow.nn.bidirectional_dynamic_rnn for output calculations. The operations and hyperparameters of models are listed in Supplementary Table 1.

For the FRSS layers, convolution kernels were rotated 180 degrees to extract reverse complementary information in the first and second layers in parallel with normal convolutions as follows. Let *W^ℓ^* a weight tensor of size M×N×H×K (M, kernel height; N, kernel width; H, input channel number; K, output channel number) in the *ℓ*th convolution layer, defined as 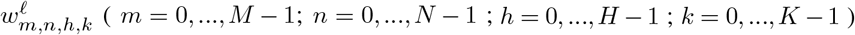. The rotated tensor 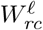 can be written as 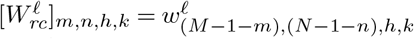. Then, the first part of the FRSS layers is

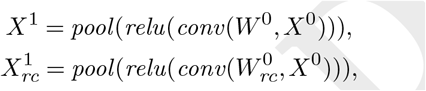

where *X^ℓ^* is a input/output tensor shape of B×S×N×H (B, mini batch number; S, sequence length; N, input width; H, input channel number) in the *ℓ*th convolution layer, 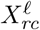 is the output of convolution with the rotated tensor, conv is the convolution operation, relu is the rectified linear unit, and *pool* is the max pooling operation. The second part of the FRSS layers is

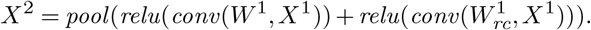

Note that the variables of 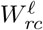 are not independent training targets, but rather are the copies of those of *W^ℓ^* .

### Training and testing models

Stochastic gradient-based optimization was used for training models. For human data, chromosomes 8 and 9 were excluded from training data. For mouse data, chromosome 2 was excluded. The excluded sequences were used to test trained models. hg19 genome was used for comparisons with previously reported AUPRCs (Fig. 2a and b). A training dataset was subdivided into minibatches containing 100 sequences. In the quick benchmark, because the fraction of positively labeled genomic windows was too small, negative samples were randomly downsampled to 25%. To accelerate the learning rate and compensate for the limited data size, each mini-batch was further divided into two sub-mini-batches, and updates were made alternately twice per sub-mini-batch (i.e., four updates per mini-batch). The loss function we used for training our models and the DeepSEA model is

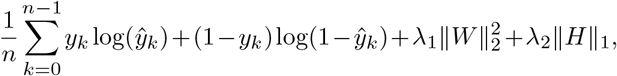

where *y_k_* is labels, 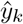 is predictions after the sigmoid operation, 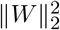 is the L2 regularization of all variables, ||*H*||_1_ is the L1 regularization of predictions before the sigmoid operation, *λ*_1_ is 5×10^-7^, and *λ*_2_ is 1×10^-8^. For the remaining models, the loss function without the regularization terms was used. The optimization algorithms RMSprop (39) and Adam (40) were used to train the DanQ model and the others, respectively. To monitor training accuracy, models were tested by every mini-batch before updating variables and then evaluated based on the F1 score 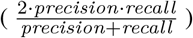 to assess temporal accuracy. When a mean of F1 scores of the last three tests exceeded a certain threshold (0.75), models were tested by three mini-batch sets that had been randomly excluded from a training dataset and saved when a higher F1 score was attained relative to the previous test.

NVIDIA TITAN X (Pascal; 3584 CUDA Cores; total amount of global memory, 12190 Mbytes; GPU Max Clock rate, 1531 MHz) was used for GPU-computing. Intel Xeon Processor E5-2640 v4 (2.40 GHz) was used for CPU-computing (see Supplementary Notes for the full specification of our machine). Total training time was dependent on the number of classes, the amount of training data, and model architecture. For example, conv4-FRSS took 3 hours to train with the subset of mm10 DNase-seq data (three classes) and 11 hours with hg38 DNase-seq (365 classes). The bottleneck was, in part, to send tensor data from the GPU to CPU for evaluation of the temporal accuracy of the training models. In previous studies, training time was reported as 85 hours in the Basset paper with NVIDIA Tesla K20m and 164 classes, and as approximately 15 days in the DanQ paper with NVIDIA Titan Z and 919 classes. However, a fair comparison of training time with previous studies is difficult owing to differences in the machines, languages, and datasets.

To test trained models, the chromosome 2 sequence for mice and chromosome 8 and 9 sequences for humans were divided into the same window size and stride with those of the training data. AUROC and AUPRC were calculated with functions in the scikit-learn library version 0.19.1 (http://scikit-learn.org/stable/). We compared the accuracy between a model trained with the full training data and one saved during training, and adopted the one with higher AUPRCs. AUPRCs of DeepSEA and DanQ in Fig. 2a and b were obtained from Supplementary Table 3 of the DeepSEA paper (https://media.nature.com/original/nature-assets/nmeth/journal/v12/n10/extref/nmeth.3547-S3.xlsx) and auc.txt of the DanQ repository (https://github.com/uci-cbcl/DanQ) and are listed in our Supplementary Table 4. Predictions presented in Fig. 2c and Supplementary Fig. 2 were visualized using the integrative genomics viewer (41).

### The class saliency extraction method

To evaluate informative sequences at the single-nucleotide level (Supplementary Fig. 4), the class saliency extraction method was performed as previously described (26) with slight modifications. Let *ŷ_i_* (*X*^0^) be the score of the class i, computed by the conv4-FRSS model for an arbitrary DNA sequence *X*^0^ of size 1000×4×1. The derivative of *ŷ_i_* (*X*^0^) with respect to *X*^0^ is

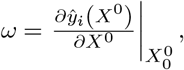

where 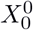 is a given DNA sequence. The derivative *ω* is a tensor of size 1000×4×1, and each element corresponds to each element of *X*^0^. The values shown in Fig. 2c and Supplementary Fig. 4a (orange) are 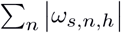 (s=0, …, 999; *n*=0, …, 3; *h*=0). To remove irrelevant values, the figures only show sequences with *S_i_* (*X*^0^) ≥ −2.0. These values represent the magnitude of the effect on the class score when nucleotide substitutions occur. Therefore, these values may be termed as influential values.

### FRiP calculation and correction

To calculate FRiPs, initially, mitochondrial and black-listed regions (https://sites.google.com/site/anshulkundaje/projects/blacklists) were removed from peak files. A module, “countReadsPerBin.CountReadsPerBin” in deeptools(42) was used to count reads in peaks, and these read counts were then divided by total reads. As a correction for read number differences, FRiP was multiplied by the fraction of genome regions covered by at least one read.

### Visualizing the inside of a trained model

Variables in each kernel were converted to a probability matrix by a softmax-like operation. Tomtom (32) version 4.12.0 and MEME motif database version 12.15 (http://meme-suite.org/meme-software/Databases/motifs/motif_databases.12.15.tgz) were used to identify closely related known transcription factor binding motifs. For visualizing kernels in the first layer of FRSS, the probability matrices were scaled by information content. Motif logos except Tomtom outputs were generated by our customized codes using the cairocffi library (https://cairocffi.readthedocs.io/en/latest/).

To visualize the connections of kernels and layers, we examined how values flow in the network, as follows. The first input sequence is a tensor of size B×1000×4×1 or 1000×4×1 (B is the minibatch number and can be ignored here). By the first convolution, in which each of 320 kernels has a 9×4 matrix, the number of output sequences is 320, and each sequence length becomes 982×1 (982×1×320). After 2×1 max pooling, the sequence length is reduced to 496×1×320. Therefore, “hidden1” in Fig. 3b and Supplementary Fig. 6 is composed of 320 dots, and each dot contains 496×1 values. Because these 320 dots are the outputs of 320 kernels, each dot has a corresponding kernel (the bottom part of Fig. 3B, but strongly connected kernels to “hidden2” are shown). From “hidden1” to “hidden2”, he number of kernels is 480, and each kernel has 320 matrices of size 9×1. Each of the 320 matrices in a kernel operates a convolution with one of the 320 outputs from the first convolution. Then, each kernel sums the 320 convolution results, which generates 480 outputs (the dots of “hidden2” in Fig. 3b) with a sequence of length 488 in each (i.e., a tensor of size 488×1×480). Namely, if a matrix of a kernel has large positive values, a corresponding output from the first convolution (a dot of “hidden1”) strongly contributes to one of 480 outputs (a dot of “hidden2”). Therefore, the sum of 9 values in a matrix of a kernel approximates the strength of a connection between two dots of “hidden1” and “hidden2”. The same algorithm was applied to the other convolution layers. The connections between the fully connected layers are simpler, as each variable represents one connection between layers. The figures present only the top 10% of the strong connections. The networks were written by our customized codes using the cairocffi library.

### The activation maximization method

Let *ŷ_i_(X*^0^) be the score of the class *i* = 0, 1, 2, computed by the conv4-FRSS model for a sequence *X*^0^. To find a sequence that is specific to the class *i*, we sought to optimize the following problem:

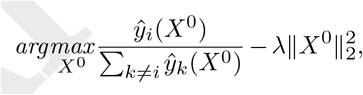

where *λ* is 5.0×10^-3^. For the common sequence (right panel in Fig. 4), the problem was:

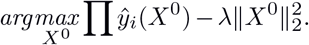

These formulae differ slightly from those reported previously (26,31) because our model used the sigmoid function for the last output, whereas the others used softmax. The optimizer Adam was used to optimize the target sequence. The initialization of variables of ***X^0^*** was set to have a mean of 0.02 and stddev of 0.02, and this was a critical factor for this optimization problem. Training with different initialization conditions sometimes did not converge, suggesting that the optimization landscape is rugged and there is a possibility that the results shown in Fig. 4 are not the optimal solution. The generated sequences were visualized with our customized codes using the cairocffi library. Tomtom (32) was used to find motifs in the generated sequences.

### Code availability

All codes used in this paper is available at https://github.com/koonimaru/DeepGMAP.

### Data availability

The IDs for data downloaded from the ENCODE website are listed in Supplementary Table 2. The data generated and/or analyzed in the current study are available in the figshare repository, https://doi.org/10.6084/m9.figshare.6728348.

## ACKNOWLEDGEMENTS

We thank Dr. Yuichiro Hara for critical comments and Dr. Mitsutaka Kadota for discussions concerning unpublished data. This work was supported in part by JSPS KAKENHI grant number 17K15132, a Special Postdoctoral Researcher Program of RIKEN, and a research grant from MEXT to the RIKEN Center for Life Science Technologies and RIKEN Center for Biosystems Dynamics Research.

